# METTL3 promotes human coronavirus replication through an interferon-independent mechanism

**DOI:** 10.64898/2026.09.18.752695

**Authors:** Ikshitaa Dinesh, Joaquim M.G.D. Fonseca, Denise Ohnezeit, Gillian Elliott, Angus C. Wilson, Hannah M. Burgess

## Abstract

N6-methyladenosine (m^6^A) is a pervasive mRNA modification that regulates RNA fate through effects on RNA-protein interactions, stability and translation. We previously showed that replication of human betacoronaviruses, OC43 (hCoV-OC43) and SARS-CoV-2, is sensitive to depletion or pharmacological inhibition of the m^6^A RNA methyltransferase METTL3, resulting in reduced viral RNA and protein accumulation. In other viral systems, such antiviral effects have been attributed to enhanced interferon (IFN) signalling and interferon-stimulated gene (ISG) induction. Here, using hCoV-OC43 we show a requirement for METTL3 that is independent of canonical IFN responses. Pharmacological inhibition of METTL3 with STM2457 failed to potentiate type I IFN signalling, global ISG expression, or the non-canonical inflammatory transcriptional programme associated with OC43 infection. Furthermore, pathogen-associated RNA sensing by RIG-I or MDA5 is not required for the antiviral effect of the STM2457. ISGs reported to be most potently antiviral against OC43 are either not significantly induced by METTL3 inhibition during infection or are not required for the antiviral activity. Nevertheless, defects in viral gene expression and progression through the viral life cycle are detectable within 6 h of STM2457 treatment and host cell transcription is dispensable for STM2457 antiviral activity. Lastly, a METTL3-directed Proteolysis Targeting Chimera (PROTAC) phenocopied STM2457, producing IFN-independent antiviral activity and ruling out off-target inhibition of viral RNA methyltransferases as a plausible explanation. Together, these findings define a direct, proviral role for METTL3 in coronavirus infection consistent with a model in which METTL3-catalysed m6A modification of viral RNA is required for efficient viral life cycle progression.

## Introduction

N6-methyladenosine (m^6^A) is the most prevalent internal base modification in eukaryotic mRNAs and can impact transcript processing, export, localisation, translation and stability by altering interactions with RNA-binding proteins and RNA structure (1, 2). It is installed by the methyltransferase METTL3, which functions as part of a larger writer complex and is predominantly nuclear localised. Nevertheless, m^6^A has been identified in viral RNAs, including those with cytoplasmic replication strategies such as the coronavirus family (3–8). Lifecycles of viruses from diverse families have now been shown to be affected both positively and negatively by perturbation of m^6^A writer complex function through genetic and pharmacologic targeting (9). It remains challenging however to differentiate whether such changes result from reduced modification of viral RNA, or indirect effects of altered host gene expression, including antiviral responses.

The replication of the large ∼30 kb positive sense RNA genomes of coronaviruses is performed by a viral RNA-dependent RNA polymerase (RdRP) and takes place in endoplasmic reticulum derived double-membrane vesicles (DMVs), which shield replication intermediates from host recognition in the cytoplasm (10, 11). Discontinuous transcription also generates nested shorter subgenomic (sg) mRNAs from which structural and accessory proteins are translated. We previously showed pharmacological targeting of METTL3 by small molecule inhibitor STM2457 impaired both SARS-CoV-2 and seasonal betacoronavirus OC43 replication in multicycle infections and the accumulation of viral RNAs (3). Inhibition of METTL3 thus provides a new potential host-directed antiviral modality, however the exact mechanism underpinning this restriction and human betacoronavirus’s requirement for METTL3 activity is not understood.

Using methylated RNA-immunoprecipitation and sequencing (meRIP-seq) and nanopore direct RNA sequencing, we and others reported evidence that coronaviral RNAs are m^6^A methylated (3–7). This raises the possibility that modified viral RNAs are regulated by m^6^A-responsive RNA-binding proteins similarly to cellular transcripts. Additionally, in vitro studies have shown that m^6^A has the potential to impact RNA-RNA interactions essential for proper viral RNA transcription and translation (12, 13). Impairment of METTL3 activity toward viral RNAs or m^6^A-mediated host gene regulation could also however influence recognition of viral RNA and the magnitude of an innate immune response. RIG-I was shown to differentially recognize m^6^A containing versus non-modified RNA ligands and viral RNAs (14, 15). Expression of multiple signalling components downstream of pathogen recognition in the interferon (IFN) pathway are also reportedly regulated by m^6^A (16–19) and the human IFN beta transcript is m^6^A-modified (20, 21), as are many interferon stimulated gene (ISG) transcripts (22). While we previously found that repression of OC43 and SARS-CoV-2 infections by STM2457 was retained when signalling downstream of the IFN receptor (IFNAR) was pharmacologically blocked (3), others have linked enhanced RIG-I recognition and a heightened antiviral host response to SARS-CoV-2’s sensitivity to METTL3 depletion (4).

Here we investigate the mechanism of restriction of seasonal human betacoronavirus OC43 by METTL3 inactivation. Interestingly, we find that the antiviral activity of METTL3 inhibition occurs independently of type I IFN signalling, RIG-I-mediated RNA sensing, and host transcription and instead our data are consistent with a model in which METTL3 directly regulates coronavirus RNA biology and viral replication. We also identify a novel host-directed antiviral strategy by demonstrating that pharmacological targeted degradation of METTL3 using a Proteolysis Targeting Chimera (PROTAC), also restricts OC43 replication independent of type I IFN signalling.

## Methods

### Cells and Viruses

MRC-5 cells were maintained in DMEM supplemented with 5% FBS and penicillin/streptomycin. HEK293T cells were maintained in DMEM supplemented with 10% FBS and penicillin/streptomycin. HCoV-OC43 was obtained from ATCC (VR-1558) and propagated in MRC-5 cells. Stocks were prepared by infecting 95% confluent T75 flask of MRC-5 cells at multiplicity of infection (MOI) of 0.1 in infection media (DMEM, 1% FBS, penicillin/streptomycin) followed by incubation for 3 days at 33°C. Supernatant was harvested and centrifuged at 1000 *g* for 5 min to remove debris. Tissue culture infectious dose 50 (TCID50) of stocks and experiment media supernatants was determined on MRC-5 cells by scoring for CPE after incubation for at least 7 d at 33°C using the Reed Muench formula. Working titre (plaque-forming units per mL) of stocks was estimated to be 0.7 TCID50/ml. Experimental infections were similarly performed in infection media (DMEM, 1% FBS, penicillin/streptomycin) and at 33°C.

### Drug and cytokine treatments

STM2457 (Cambridge Bioscience) was reconstituted in DMSO to a 10 mM stock concentration and used at a final concentration of 30 µM. Single use aliquots were stored at -80°C. Recombinant human IFN-β (Proteintech HZ-1298) was reconstituted in water to 200 µg/ml from a 10 µg lyophilised stock and stored at -80°C. JAK inhibitor Pyridone 6 (Millipore-Sigma 420099) was reconstituted in DMSO to a stock concentration of 10 mM, stored at -20°C and used at a final concentration of 10 µM. Actinomycin D was reconstituted in DMSO to 10 mg/ml, aliquots stored at -20°C and used at a final concentration of 15 µM. WD6305 (Insight Biotechnology) was reconstituted in DMSO to a 1 mM stock, stored at -80°C and used at the indicated concentrations.

### Plasmids and transfection

Plasmid transfections were performed using Lipofectamine™ 2000 (Life Technologies) using 1 μg total plasmid DNA per ml. Plasmids encoding C-terminally FLAG-tagged OC43 Nsp14 and Nsp16 (strain VR-1558) were generated in a pcDNA3.1 backbone by GeneArt (Thermo Fisher Scientific). The OC43 N-FLAG plasmid was generated by transferring the OC43 nucleoprotein (N) coding sequence from pGBW-m4134899 (gift of Ginkgo Bioworks & Benjie Chen, Addgene #151902) into pFLAG-CMV-5a.

### RNA interference

Cells were seeded and transfected the next day using 3 µL/mL Lipofectamine RNAiMax (Life Technologies) at a final siRNA concentration of 20 nM. Three days after transfection, cells were infected or treated as indicated. The following siRNAs were used: Allstars negative control siRNA (Qiagen, #1027281) DDX54/RIG-I (Sigma SASI_HS01_00047980), IFIH1/MDA5 (Sigma SASI_Hs01_00171929), OAS1 (Sigma SASI_Hs01_00154844), OAS2 (Sigma SASI_Hs01_00054392) and OAS3 (Sigma SASI_Hs01_00104798).

### RT-qPCR analysis

Cells were lysed and RNA was isolated using a Qiagen RNeasy Plus mini kit following manufacturer’s protocol. RNA was reverse transcribed using qScript Ultra supermix (Quanta). Quantitative PCR (qPCR) reactions were conducted using Takyon No ROX MasterMix dTTP Blue (Eurogentec) and a BioRad CFX connect quantitative PCR system. For each biological replicate, technical duplicates were conducted. Fold changes in gene expression relative to GAPDH were calculated using the ΔΔCT method. Statistical analyses on RT-qPCR data were performed using GraphPad Prism, using tests indicated. Primer sequences are as follows: 18S (Fwd AGGAATTGACGGAAGGGCACC; Rev TTATCGGAATTAACCAGACAAATCG) GAPDH (Fwd TCTTTTGCGTCGCCAGCCGA; Rev ACCAGGCGCCCAATACGACC); OC43 genomic ORF1ab (Fwd TCCTACTTGGAGTCAGGAACT, Rev GTAGCAGCACTCTGCAATCT); OC43 sgN (Fwd TCCCGCTTCACTGATCTCTT, Rev TTTGCTTGGGTTGAGCTCTT); IFNB (Fwd GAAAGAAGATTTCACCAGGG, Rev CCTTCAGGTAATGCAGAATC); MX1 (Fwd GGCTGTTTACCAGACTCCGACA, Rev CACAAAGCCTGGCAGCTCTCTA); ISG15 (Fwd AGATCACCCAGAAGATCG, Rev TGTTATTCCTCACCAGGATG); DDX58 (RIG-I) (Fwd GGTATAGAGTTACAGGCATTTC, Rev TTGTTTACTAGTGTTGTGGC); IFIH1 (MDA5) (Fwd GATTAAGTGGTGATACCCAAC, Rev GTCTGACAATTGAACACCAG) CXCL3 (Fwd CGCCCAAACCGAAGTCATAG, Rev GCTCCCCTTGTTCAGTATCTTTT); CXCL1 (Fwd CGGAAAGCTTGCCTCAATCC, Rev GGTCAGTTGGATTTGTCACTGTT); IL1A (Fwd GCGTTTGAGTCAGCAAAGAAGT, IL1A Rev CAGAGACAGATGATCAATGGAGGA) IL6 (Fwd GCAGAAAAAGGCAAAGAATC, Rev CTACATTTGCCGAAGAGC); CXCL8 (IL8) Fwd GCGCCAACACAGAAATTATTGTAAA, Rev TGAATTCTCAGCCCTCTTCAAAAA); CXCL10 (Fwd AAAGCAGTTAGCAAGGAAAG, Rev TCATTGGTCACCTTTTAGTG); CXCL11 (Fwd GCTACAGTTGTTCAAGGCTTC, Rev AGGCTTTCTCAATATCTGCCACT); LY6E (Fwd TGCCGGCATTGGGAATCTC, Rev ACATTGACGCCTTCTGGGAT); TNFRSF10A (Fwd TTGTTGCATCGGCTCAGGTTG, Rev TCTCGTTGTGAGCATTGTCCT); SCARB1 (Fwd GACCAATCTGTTGGAGACCCT, Rev GCACCTGGGACCACTCTATG); NCOA7 (Fwd AGAGCCACACTTCTCACTGC, Rev TCGTCAGGTCTGGCACAATAG); ANKFY1 (Fwd AGCAGCGAGTCCTTCATCAG, Rev TTGTGAGCACTGATGTGCCT); UNC93B1 (Fwd TACAACGAGGAGGAGGAGGA, Rev AGGTAGACGCCGTAGGTGA); CTSS (Fwd TGGATCACCACTGGCATCTCTG, Rev GCTCCAGGTTGTGAAGCATCA); ETV6 (Fwd AGTGTAGCATTAAGCAGGAACGA; Rev ATGAAGTGGCGTCGAGGAAG); OAS1 ( Fwd AGGAAAGGTGCTTCCGAGGTA, Rev GGACTGAGGAAGACAACCAGGT); OAS2 (Fwd ATTGGATGGTCAACTACAAC, Rev GCAGGTAACTCATTATCCAG) OAS3 (Fwd TGCTGCCAGCCTTTGACGCC, Rev TCGCCCGCATTGCTGTAGCTG).

### Immunoblotting

Protein lysates were collected in NuPAGE LDS Sample Buffer (Invitrogen, NP0007) and subject to SDS-PAGE. Following transfer to nitrocellulose, membranes were incubated with primary antibodies: GAPDH (Cell signalling 2118; 1:1000), MDA5 (Proteintech 21775-1-AP; 1:5000), METTL3 (Abcam ab195352; 1:1000), METTL14 (Sigma HPA038002; 1:1000), OC43 nucleocapsid (Millipore 541-8F; 1:4000) and RIG-I (Proteintech 20566:1:2000). Secondary antibodies used were as follows: goat anti-rabbit IRDye 800CW (Licor 926-3211), goat anti-rabbit 680RD (Licor 926-68071), and goat anti-mouse IRDye 800CW (Licor 926-68070). Blots were imaged and processed on a Licor Odyssey system and Licor Image Studio software. All immunoblots presented are representative of at least 2 independent experiments.

### Direct RNA sequencing

RNA was isolated from MRC-5 cells infected with OC43 at an MOI of 3 for 24 h in the presence of DMSO or STM2457 using TRIzol (Invitrogen) reagent, as per the manufacturer’s instructions. RNA was precipitated using ethanol and sodium acetate and pellets were washed twice with 75% ethanol. Poly(A) RNA enrichment was conducted using the Dynabeads™ mRNA Purification Kit (Invitrogen), following the manufacturer’s instructions. 300 ng of poly(A) RNA were used as input for library preparation using ONT’s standard DRS SQK-RNA004 protocol. Libraries were loaded onto individual PromethION RNA flow cells (ONT) and sequenced on a PromethION 2 solo device, acquiring 4M reads per sample. Basecalling was performed using Dorado v0.9.0 (ONT) in super accuracy mode with modification detection enabled for m6A (DRACH context). Human transcriptome alignment and modification mapping was performed as previously described (23). For viral alignment, reads were aligned against the HCoV-OC43 reference genome (NC_006213.1) using minimap2 v2.28 [minimap2 -ax splice -k 8 -w 3 -g 30000 -G 30000 -C0 -un -no-end-flt -splice-flank=no], or against HCoV-OC43 transcripts [minimap2 -ax map-ont -L -p 0.99 -y --secondary=no]. Using SAMtools vl.16, genome-and transcriptome-aligned SAM files were converted to BAM files retaining only primary alignments [samtools view -b -F2308 and samtools view -b -F2324, respectively], with subsequent sorting [samtools sort] and indexing [samtools index]. Plots were produced by ggplot2 using R version 4.4.0.

### Data availability

Sequencing data sets associated with this study have been deposited as fast5 files at the European Nucleotide Archive (ENA) under the project accession PRJEB126705.

### Statistical Analyses

Statistical analyses were performed using Graphpad Prism. Where samples were normalised to a control value set to 1, a one sample t-test was applied. In all figures ns is not significant, * p < 0.05, ** p < 0.01, *** p < 0.001 and **** p<0.0001.

## Results

### METTL3 inhibition reduces OC43 RNA within 6 hours

We previously found STM2457 reduced OC43 gene expression in synchronised single cycle infections of MRC-5 cells within 24 hours (3). To determine the earliest time when the antiviral impact of STM2457 on viral RNA accumulation can be detected, we performed a synchronised time course of infection (Fig.1A). MRC-5 cells were infected at high multiplicity (MOI: 3) to exclude any effects on viral entry and to minimize the impact on host gene expression, and cells were treated with STM2457 (30 μM) or a vehicle control (DMSO) only after viral adsorption (1 h). Total RNA was harvested at 3 h intervals up to 12 hours post infection (hpi) and at 24 hpi. RT-qPCR was performed to measure genomic RNA, using primers that amplify a product within the ORF1a region (Fig. 1A), and the abundant subgenomic (sg) mRNA encoding the nucleocapsid (N) protein (Fig. 1B). Both species of viral RNA were detectable within 3 h of infection, and abundance scaled exponentially up to 24 h (Fig. 1A, B, upper panels). Normalising viral RNA expression in STM2457 treated conditions to DMSO treatment (Fig. 1A, B lower panels), revealed that METTL3 inhibition leads to a detectable reduction in genomic and subgenomic viral RNA accumulation within 6 hours of infection, corresponding to 5 h of STM2457 treatment. These data are consistent with either a requirement for METTL3 activity for an early viral process, such as transcription or translation, or for the control of a host gene or genes that potently regulate infection within this time frame.

**Figure 1.**
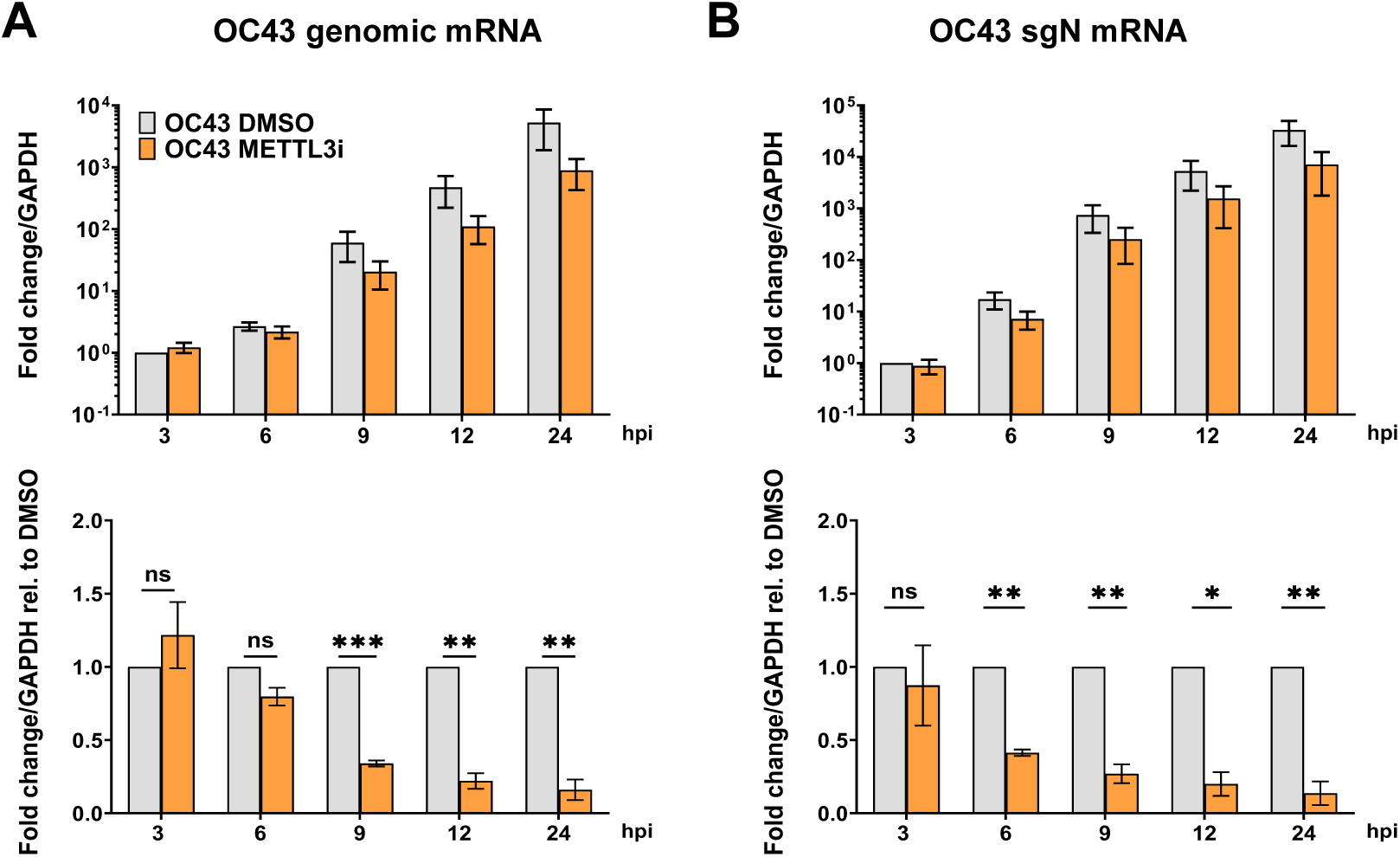
METTL3 inhibition reduces viral RNA levels within hours of infection. MRC-5 cells were infected with HCoV-OC43 at an MOI of 3 and after 1 h of viral absorption treated with 30 µM STM2457 or DMSO. (A) RNA was harvested at the time indicated and analysed by RT-qPCR using primers specific for viral genomic (ORF1ab) or sub-genomic (N) mRNA. Mean fold changes relative to GAPDH ± SEM are plotted, n ≥ 3. In the upper panel expression is normalised to DMSO-treated samples at 3 hpi. In the lower panel expression is normalised to DMSO-treated samples at each timepoint. Statistical significance was tested on timepoint-normalised values by one sample t-test.

### OC43 is acutely sensitive to type 1 IFN

Dysregulation of the type 1 interferon antiviral host response has been implicated in the control of several DNA and RNA viruses by METTL3 through a range of mechanisms, including the stability of the IFNB transcript (20, 21), stability of interferon stimulated gene (ISG) mRNAs (22) and differential recognition of modified viral RNAs by cellular pathogen recognition receptors (14, 24, 25). As such, STM2457 treatment could conceivably act by enhancing the antiviral response to OC43. To test whether OC43 can be restricted by type-I IFN, MRC-5 cells were treated with a range of concentrations of human IFN beta (IFN-β) for 24 h and subsequently infected with OC43 at low multiplicity (MOI: 0.001). At 48 hpi released virus titres were quantified by TCID50 assay (Fig. 2A). Expression of ISGs immediately prior to virus infection (24 h IFN-β treatment) was also validated by RT-qPCR for MX1 and ISG15 (Fig. 2B). OC43 replication was significantly and potently (>3 log_10_) reduced by pretreatment with 0.1-10 ng/ml IFN-β. Accordingly, the same concentrations of IFN-β resulted in massive upregulation of MX1 and ISG15, with ∼300-fold induction of MX1 mRNA corresponding with the IFN mediated restriction of OC43 at 0.1 ng/ml. While 0.01 ng/ml IFN-β treatment upregulated MX1 97-fold and ISG15 18-fold, this did not reach statistical significance and was not associated with OC43 replication restriction.

**Figure 2.**
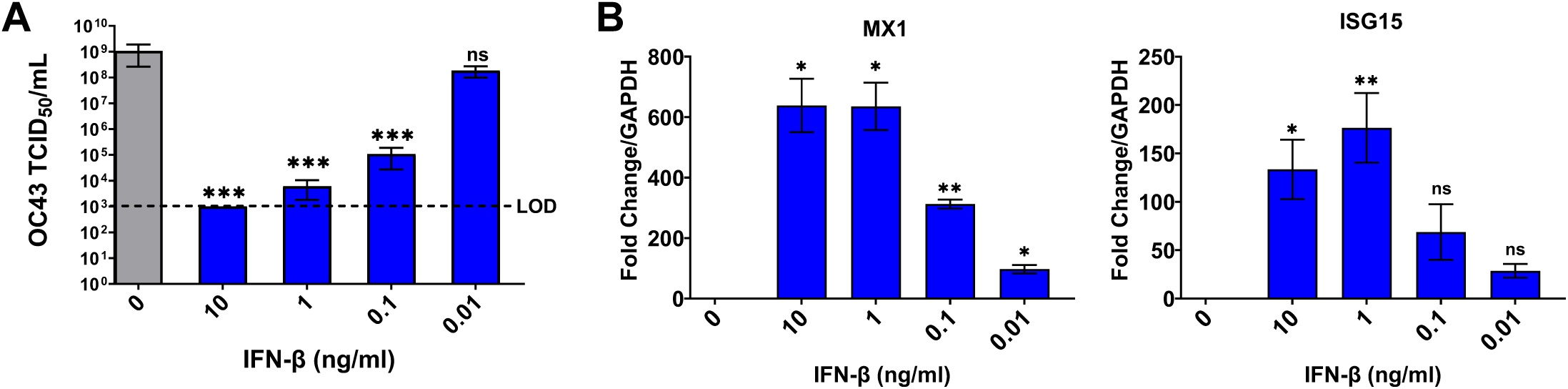
OC43 is acutely sensitive to type 1 IFN. MRC-5 cells were pre-treated with indicated concentrations of IFN-β for 24 h. (A) Pre-treated cells were infected with OC43 at MOI 0.001 and media supernatant collected at 48 hpi and yield of infectious virus determined by TCID50 assay. Dashed line indicates assay limit of detection (LOD). Mean ± SEM is plotted, n ≥ 3. Statistical significance was tested on transformed values in comparison to control treated samples by ordinary one-way ANOVA with Dunnett’s multiple comparison correction. (B) RNA was harvested from uninfected pre-treated cells and analysed by RT-qPCR for ISG expression, shown as mean ± SEM relative to GAPDH and normalised to untreated control (n ≥ 3). Statistical significance was tested by one sample t-test.

### STM2457 does not enhance a global canonical ISG or inflammatory response to OC43 infection

Since IFN-β pre-treatment can impact OC43 replication, we next asked if STM2457 leads to increased expression of IFN-β and canonical ISGs during infection. MRC-5 cells were infected with OC43 at high MOI (3) and either DMSO or STM2457 was added to the cultures following viral adsorption. RNA was collected at 24 hpi and RT-qPCR performed (Fig. 3A). First viral RNA levels were assessed and consistent with the previous time course samples, showed a reduction upon STM2457 treatment. Interestingly, IFNB mRNA levels were only modestly upregulated by infection (< 3-fold) and not further enhanced by STM2457 treatment (Fig. 3A). While MX1 was upregulated by infection (10-fold), this was much less than the induction associated with IFN-β treatment concentrations that restricted OC43 replication (Fig. 2), and importantly, this was not enhanced by STM2457 treatment. ISGs DDX58 (RIG-I) and IFIH1 (MDA5) showed a similar pattern while ISG15 was not detectably induced by OC43. Recent transcriptomic analysis of OC43 infection of MRC-5 cells demonstrated that, consistent with our RT-qPCR data, the virus largely evades a broad induction of canonical ISGs, instead eliciting a distinct inflammatory cytokine/chemokine signature, including upregulation of IL1A, IL6, CXCL1, CXCL3, CXCL8, CXCL10 and CXCL11 (26). Further RT-qPCR analysis confirmed moderate upregulation of these genes by OC43 infection, but in no case did STM2457 treatment enhance their expression during infection. Indeed, for DDX58 and several other chemokines (IL6, CXCL2, CXCL3), STM2457 resulted in a significant reduction in expression during infection (Fig. 3A). Together these data demonstrate that the canonical type I IFN response or the broad inflammatory gene signature induced by OC43 infection are not amplified by STM2457.

**Figure 3.**
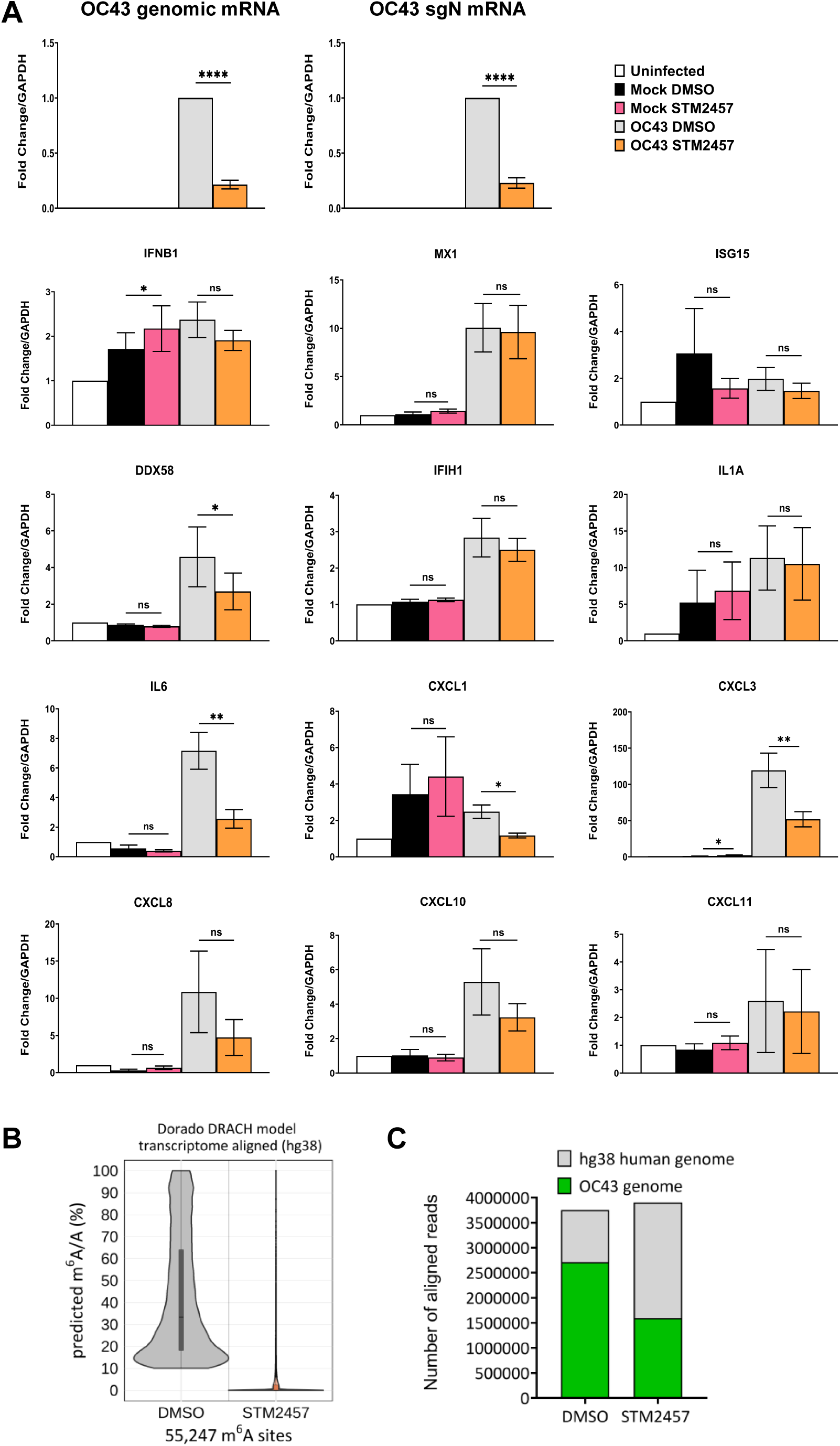
STM2457 does not enhance a global ISG or inflammatory response to OC43. MRC-5 cells were infected with OC43 at MOI of 3 and treated with STM2457 (30 µM) or DMSO control as in Fig. 1. RNA was collected at 24 hpi. (A) RNA was subject to RT-qPCR for the indicated viral RNAs, host cytokines, chemokines and ISGs. Mean fold changes ± SEM are plotted relative to GAPDH and normalised to untreated uninfected control, or for viral RNAs normalized to DMSO-treated infected samples, n ≥ 3. Statistical significance was tested by two-tailed paired t-test. (B) RNA was subject to direct RNA sequencing and the modification-aware basecaller Dorado (DRACH model) was applied to detect m6A sites over the human transcriptome. Predicted m^6^A/A proportions are plotted for high-confidence m^6^A sites detected in both DMSO and STM2457-treated cells. (C) Total numbers of DRS-captured reads aligned to both the human (hg38) and OC43 genome are shown from DMSO-or STM2457-treated cells.

Though we detected a consistent suppression on viral RNA accumulation by STM2457 treatment across our experiments, our failure to detect METTL3-dependent enhancement in innate immune responses contrasts with other studies (4, 7). To ensure STM2457 effectively blocks the installation of m^6^A on host mRNAs in our infection conditions we performed nanopore direct RNA sequencing (DRS) on total RNA isolated from MRC-5 cells infected with OC43 and treated as in Fig. 3A using the new Oxford Nanopore Technology (ONT) SQK-RNA004 chemistry. The ONT Dorado modification-aware basecaller (v0.9.0) was used to identify and quantify the predicted m^6^A/A proportions within a DRACH consensus sequence context on the host transcriptome. We previously established that stringent filtering for Dorado-predicted m^6^A modified sites with a modification probability of 98% and a predicted m^6^A/A proportion score of ≥ 10% provides accurate site identification and minimizes false positive calls (23). Under such filtering, 55,247 high confidence m^6^A-modified sites on cellular polyadenylated RNAs were identified in our samples (Fig. 3B). Stoichiometry of modification varied widely, with many (> 30%) adenosine sites modified on 10-20% of molecules, and comparatively fewer modified consistently, in line with our previous analyses of human fibroblasts with this methodology (23) and other quantitative m^6^A mapping studies (27, 28). STM2457 treatment led to a near complete loss of m^6^A on host transcripts, with median predicted m^6^A/A proportions dropping from 33% in DMSO-treated cells to 0% in STM2457-treated cells. The effect of STM2457 treatment on total viral RNA accumulation was evident in much reduced reads mapping to the OC43 genome (Fig. 3C), consistent with our RT-qPCR analysis. Thus, failure to detect an enhanced host immune and inflammatory response upon STM2457 treatment during OC43 infection is not due to incomplete inhibition of METTL3-mediated m^6^A installation.

### STM2457 anti-coronaviral activity does not require RIG-I or MDA5

A subset of ISGs can be induced directly by IRF3, without requiring IFN expression and IFNAR activation (29). RIG-I like receptor (RLR) binding to viral RNA has also been reported to negatively regulate RNA viruses without requiring MAVS signalling or antiviral gene expression (30, 31). We therefore tested if either cytosolic RNA sensor RIG-I or MDA5 are required for the antiviral activity of STM2457 against OC43. Each gene was targeted for depletion in MRC-5 cells by specific siRNAs, alongside a non-silencing control siRNA. After 3 days to allow target protein turnover, cells were infected at MOI of 3 in the presence of STM2457 or DMSO and lysates collected at 24 hpi. Immunoblotting confirmed efficient knockdown of RIG-I and MDA5 (Fig. 4A). Consistent with our previous results (3), nucleocapsid protein levels in control cells were reduced by STM2457 treatment to 47% compared to control. In RIG-I and MDA5 knockdown cells, a similar effect was observed, with a reduction in nucleocapsid protein to 37% and 50% compared to DMSO-treatment, respectively (Fig. 4B). We next tested whether any difference in STM2457 antiviral efficacy could be detected in RIG-I and MDA5 knockdown cells by measuring released virus titres after a low multiplicity infection. Here the suppressive effects of STM2457 treatment were very similar in control and knockdown cells, with STM2457 treatment resulting in a statistically significant reduction in titres in all conditions (Fig. 4C). These data demonstrate that the restriction of OC43 productive replication by METTL3 inhibition does not depend on either the signalling or non-canonical roles of RIG-I or MDA5.

**Figure 4.**
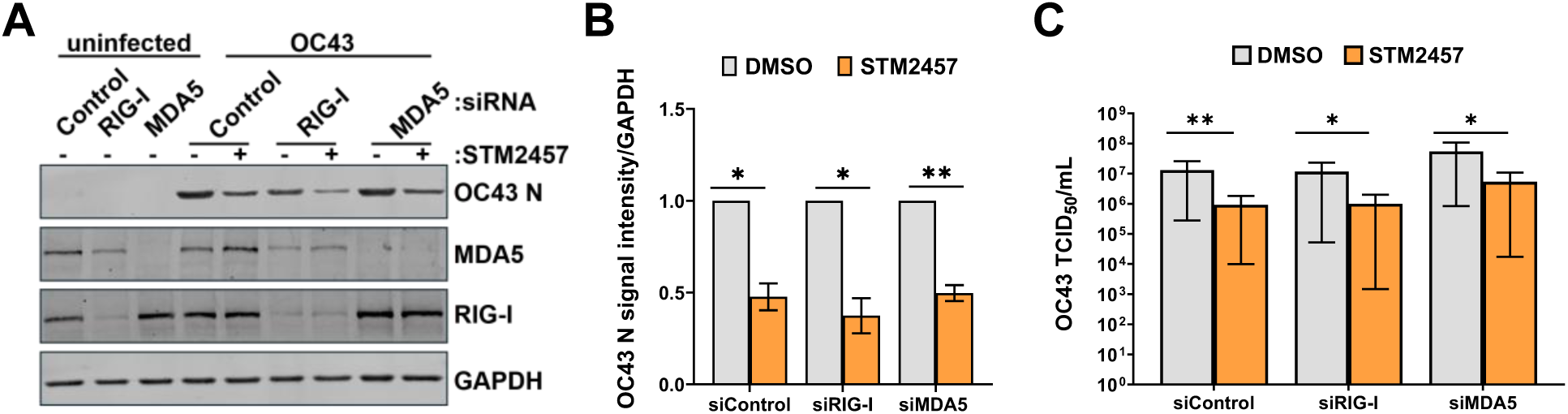
STM2457 anti-coronaviral activity does not require RIG-I or MDA5. MRC-5 cells were transfected with 20 nm siRNAs targeting RIG-I, MDA5 or a non-silencing control for 72 h. Cells were infected with OC43 at MOI 3 (A, B) or 0.001 (C) and treated with DMSO or STM2457 (30 µM). (A) Protein lysates were collected at 24 hpi and subject to immunoblotting for the indicated proteins. (B) Immunoblot signal for OC43 nucleocapsid protein was quantified and plotted as mean ± SEM (n=3), normalised to loading control GAPDH, and shown relative to DMSO for each siRNA. Statistical significance was tested by one sample t-test. (C) Released infectious viral titres were established by TCID50 assay and plotted as mean ± SEM (n=3). Statistical significance was tested on transformed values by two-tailed paired t-test.

### Dysregulation of specific ISGs directly implicated in restriction of betacoronaviruses does not explain STM2457 antiviral activity

Though hundreds of genes are upregulated by interferon, distinct subsets provide the antiviral defence against specific viruses (32). A recent overexpression study defined those ISGs most potently inhibitory for OC43 (33), following similar studies in SARS-CoV-2 (34–36). We therefore tested whether gene specific regulation by METTL3 of ISGs that selectively regulate OC43 could be involved. We used RT-qPCR to examine the expression of 10 of the ISGs most strongly implicated in control of OC43 or SARS-CoV-2 after a high multiplicity infection of MRC-5 cells in the presence of DMSO or STM2457 (Fig. 5). Because OAS1 and OAS2 were previously implicated as particularly potent inhibitors of SARS-CoV-2 and OC43, respectively, we measured the relative expression of all three human 2′–5′ oligoadenylate synthetase (OAS) genes. For most genes examined (LY6E, TNFRSF10A, SCARB2, ETV6, CTSS, UNC93B1, ANKFY1, OAS3), their expression was neither significantly upregulated by OC43, nor enhanced by STM2457 treatment during infection (Fig. 5). Similar to MX1 (Fig. 3A), NCOA7 was moderately induced ∼5-fold by OC43, but this was reduced by STM2457 treatment. Interestingly, OAS1 expression was also moderately induced by OC43 infection (17-fold) and this was increased slightly to 26-fold in the presence of STM2457. Notably, while OAS2 expression was only induced 3-fold by infection, this was increased to 6.5-fold in STM2457-treated infected cells. Though the difference in OAS1 and OAS2 expression between DMSO and STM2457-treated infected conditions did not reach statistical significance, likely due to normalisation to low and variable basal expression, as these genes are linked to coronavirus restriction and could function redundantly, we next functionally investigated the OAS gene family.

**Figure 5.**
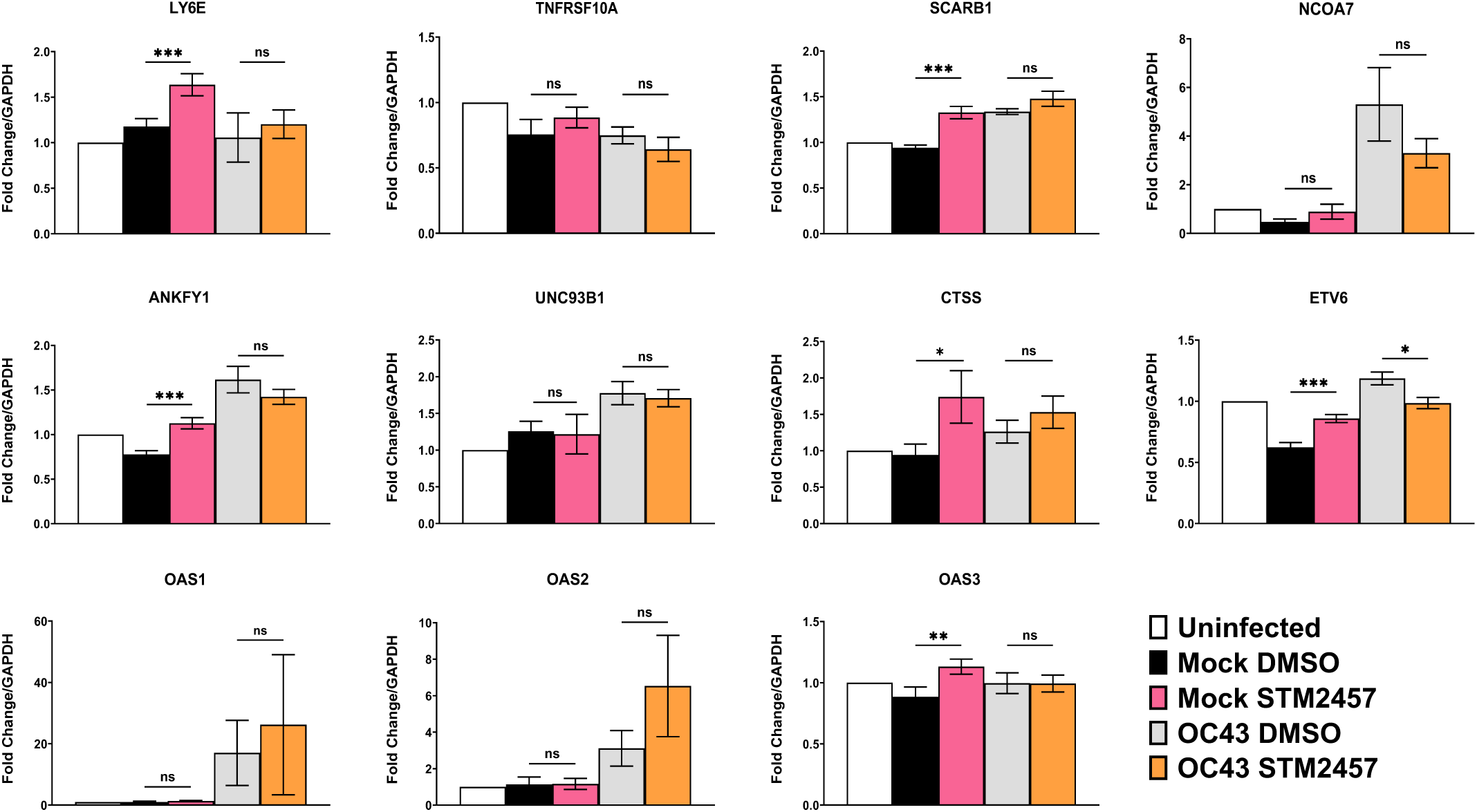
Expression of ISGs that restrict hCoVs. RNA samples in Fig. 3 were analysed further by RT-qPCR for ISGs implicated as CoV-specific. Mean fold changes ± SEM, relative to GAPDH and normalised to untreated uninfected control are plotted (n ≥ 3). Statistical significance was tested by two-tailed paired t-test.

To establish whether OAS genes contributed to the restriction of OC43 by type 1 IFN, we transfected MRC-5 cells with siRNAs targeting OAS1, OAS2 and OAS3 or the same final concentration of a non-silencing control siRNA and after 24 h treated cells with 0.1 ng/ul IFN for 24 h or left them untreated. This concentration was chosen to allow a moderate induction of ISGs and still permit OC43 replication (Fig. 2). Cells were then infected at MOI 0.001 and at 48 hpi released viral titres measured by TCID50 assay (Fig. 6A). Knockdown efficiency was measured by RT-qPCR, which demonstrated that each target transcript was reduced by more than 80% (Fig. 6B). Knockdown of OAS genes did not increase titres of virus from cells which were not pre-treated with IFN (Fig. 6A). In IFN treated cells, however, knockdown of OAS genes increased released viral titres ∼30-fold (Fig. 6A), confirming that OAS genes contribute to the restriction of OC43 by type 1 IFN. To test whether the regulation of OAS genes following treatment with the METTL3 inhibitor in OC43-infected cells could account for its antiviral activity, we tested the ability of STM2457 to repress viral protein production in control and OAS1/2/3-targeting siRNA transfected cells (Fig. 6C). Cells were transfected for 72 hours then infected at MOI of 3 in the presence of STM2457 or DMSO and lysates collected at 24 hpi for immunoblotting. In both conditions, STM2457 resulted in a very similar reduction in nucleocapsid expression.

**Figure 6.**
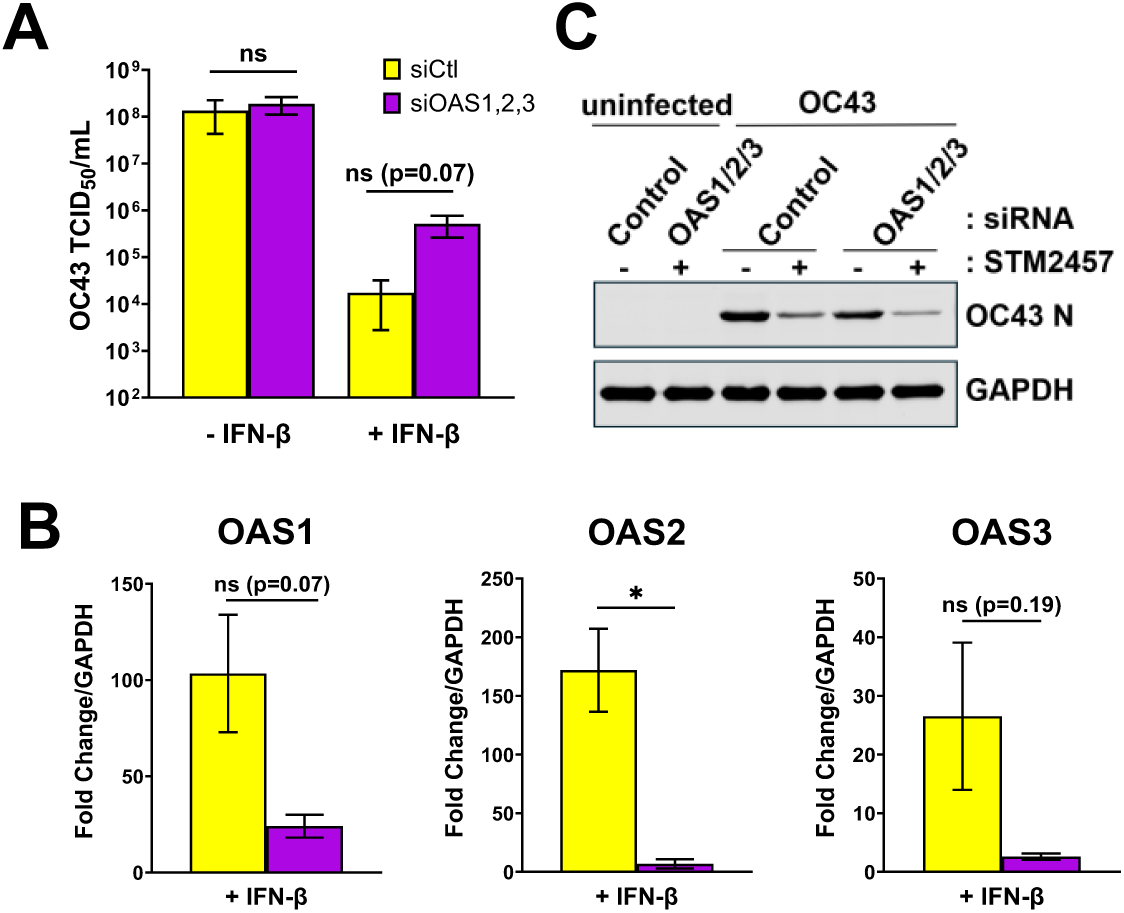
OASes are not required for METTL3 inhibition of OC43. (A) MRC-5 cells were transfected with 20 nm siRNAs targeting OAS1, OAS2, and OAS3 or 60 nM non-silencing control siRNA for 24 h before 48 h treatment with IFN-β at 0.1 ng/ml for 24 hpi or a media change. Cells were then infected with OC43 at MOI 0.001 for 48 h and viral supernatants collected and analysed by TCID50 assay and plotted as mean ± SEM (n=3). (B) Knockdown of each OAS in IFN-β-treated samples was validated by RT-qPCR at 72 hours post-transfection. Mean fold change ± SEM (n=3) was plotted normalised to siControl -IFN-β samples. Statistical significance was tested on transformed values by two-tailed paired t-test (A,B). (C) MRC-5 cells were similarly transfected with 20 nm siRNAs targeting OAS1, OAS2, and OAS3 or 60 nM non-silencing control siRNA for 72 h before infection with OC43 at MOI 3 in the presence of STM2457 (30 µM) or DMSO. Protein lysates were collected at 24 hpi and subject to immunoblotting for the indicated proteins.

Overall, we find that most ISGs that are potently antiviral for OC43 are not selectively upregulated by STM2457, and that OAS dysregulation is not required for the antiviral activity of STM2457.

### Ongoing host gene expression is dispensable for the antiviral activity of STM2457

Several genome wide studies have revealed the potential for constitutively expressed host genes to restrict replication of viruses, including SARS-CoV-2 and OC43 (37–41). Given that transcripts from around 7,000 human genes are m^6^A modified to some extent (42, 43), we next considered whether STM2457 could act indirectly by shaping the host transcriptome to antagonize OC43 replication. We first tested whether pretreatment of cells with STM2457 24 h prior to OC43 infection altered viral RNA accumulation at early time points (Fig. 7A). Cells were infected at high multiplicity (MOI 3) and total RNA was collected at 3 h intervals up to 12 hpi and at 24 hpi, similar to Fig. 1. During and after infection no STM2457 was included, thus excluding direct effects on viral RNA. In both non-treated and STM2457-pretreated cells genomic and subgenomic RNAs accumulated normally (Fig. 7A, upper panel). Directly comparing the impact of STM2457 pretreatment with DMSO pretreatment revealed no deficit in viral RNA levels, even at the earliest time following STM2457 withdrawal, 3 hpi, where a small but statistically significant increase in genomic RNA was detected after STM2457 pretreatment (Fig. 7A, lower panel). These data imply that STM2457 does not induce changes in the host transcriptome that render cells less permissive to OC43 replication.

**Figure 7.**
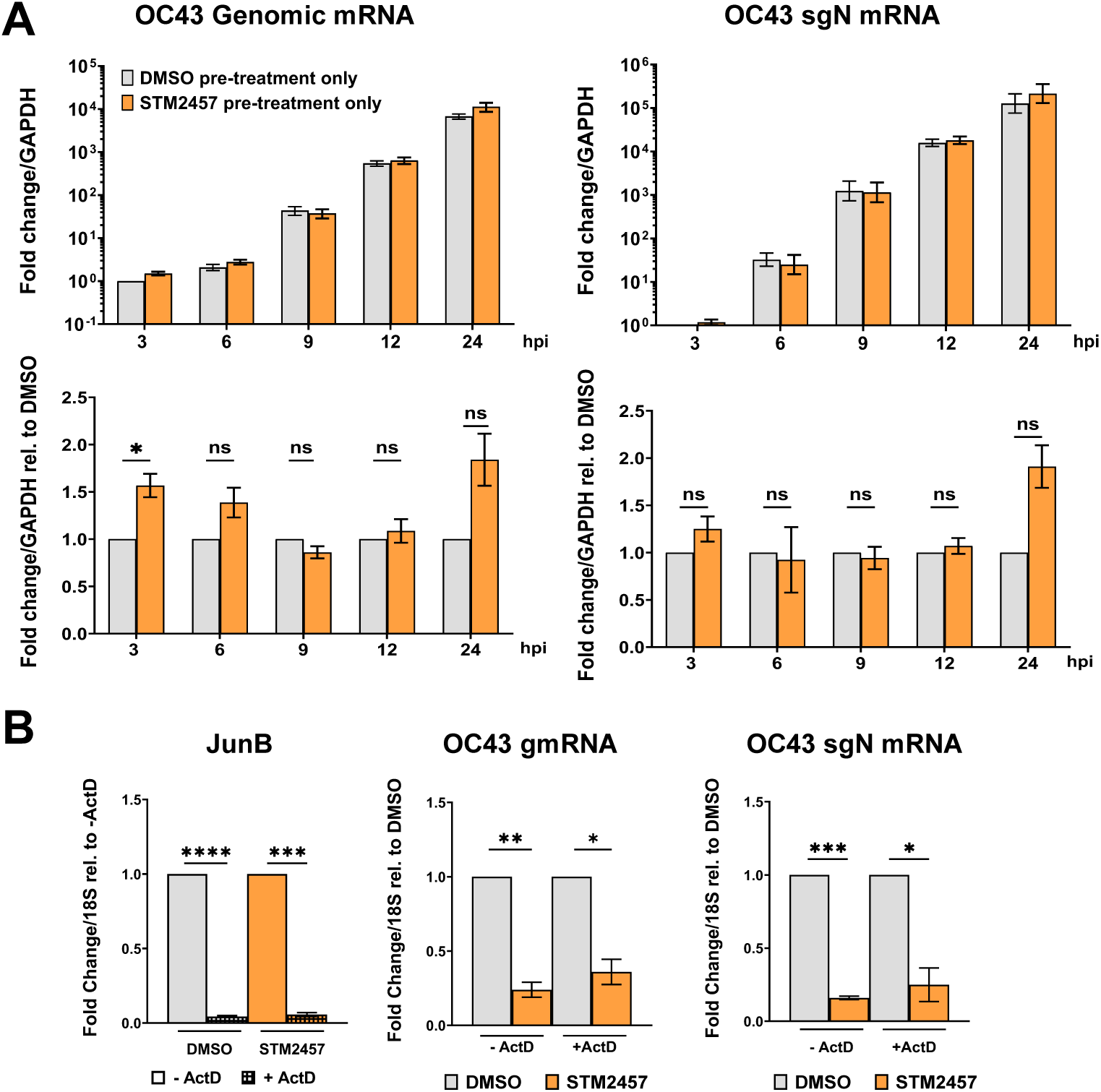
Antiviral activity of STM2457 does not require host gene expression. (A) MRC-5 cells were pre-treated with STM2457 (30 µM) or DMSO for 24 h and subsequently infected at MOI 3 without any additional treatments. RNA was collected at the indicated times post-infection and subject to RT-qPCR for viral RNAs. In the upper panel expression is normalised to DMSO-treated samples at 3 hpi. In the lower panel expression is normalised to DMSO-treated samples at each timepoint. Mean ± SEM are plotted, n=3. Statistical significance was tested on timepoint-normalised values by one sample t-test. (B) MRC-5 cells were infected with OC43 (MOI 3) in the presence or absence of 15 µM Actinomycin D and either STM2457 (30 µM) or DMSO. RNA was collected at 9 hpi for RT-qPCR analysis, with fold changes in mRNA expression normalised to 18S and plotted relative to DMSO treatment (OC43 gmRNA and sgN mRNA) or no Actinomycin D control (JunB). Mean ± SEM are shown, n=3. Statistical significance was tested by one sample t-test.

To directly test whether modification of host RNAs contributes to the antiviral activity of STM2457 we sought to isolate effects on host versus viral transcription. Modification of cellular mRNAs by METTL3 within the m^6^A writer complex is believed to largely occur co-transcriptionally (44–46). We therefore tested the ability of STM2457 to restrict viral RNA accumulation when host transcription was inhibited with actinomycin D (ActD). Coronavirus RNA synthesis is entirely achieved by a viral RNA-dependent RNA polymerase which is unaffected by ActD (47, 48). Cells were infected with OC43 (MOI 3) and treated with DMSO or STM2457 in the presence or absence of ActD (Fig. 7B). To verify ActD treatment effectively blocked host transcription RT-qPCR was performed to detect *JunB* mRNA, which has a short half-life (49). Accordingly, *JunB* levels were reduced by >96% by ActD treatment confirming its efficacy. Importantly, STM2457 remained effective in suppressing OC43 RNA accumulation in ActD treated cells and reduced both viral genomic and subgenomic RNAs to similar levels regardless of ActD treatment (Fig 7B). Together these data demonstrate that host gene expression, and thus co-transcriptional installation of m^6^A on host transcripts, is not required for STM2457’s antiviral activity toward OC43.

### A METTL3-targeting PROTAC has an IFN-independent antiviral effect

Proteolysis Targeting Chimeras, or PROTACs, are an emerging class of therapeutics that mediate proteasomal degradation of a target protein through recruitment to an E3 ubiquitin ligase. Their potential to be deployed as antivirals is only now being explored, with PROTACs aimed at both viral proteins and host dependency factors under investigation (50). Advantages over small-molecule inhibitors include prolonged efficacy, as restoration of target protein abundance requires time following degradation, and activity at sub-stoichiometric concentrations, since sustained target binding is not required.

We previously showed that siRNA knockdown of METTL3 72 h prior to infection reduced subsequent SARS-CoV-2 and OC43 replication (3) and next aimed to test whether acute depletion using a novel METTL3-targeting PROTAC, WD6305 (51), could achieve the same effect. To establish the shortest treatment time required for substantial target reduction, we first characterized METTL3 protein levels using immunoblotting following treatment with 0.5 – 5 µM WD6305 for 2 or 4 hours (Fig. 8A). At least 4 hours of WD6305 treatment was required to reduce METTL3 by more than 80%, with 4 h treatment at 1 µM yielding similar depletion (88%) to higher concentrations. Thus, treatment of cells with 1 µM WD6305 for 4 h prior to, during and after infection was used for subsequent experiments in MRC-5 cells.

**Figure 8.**
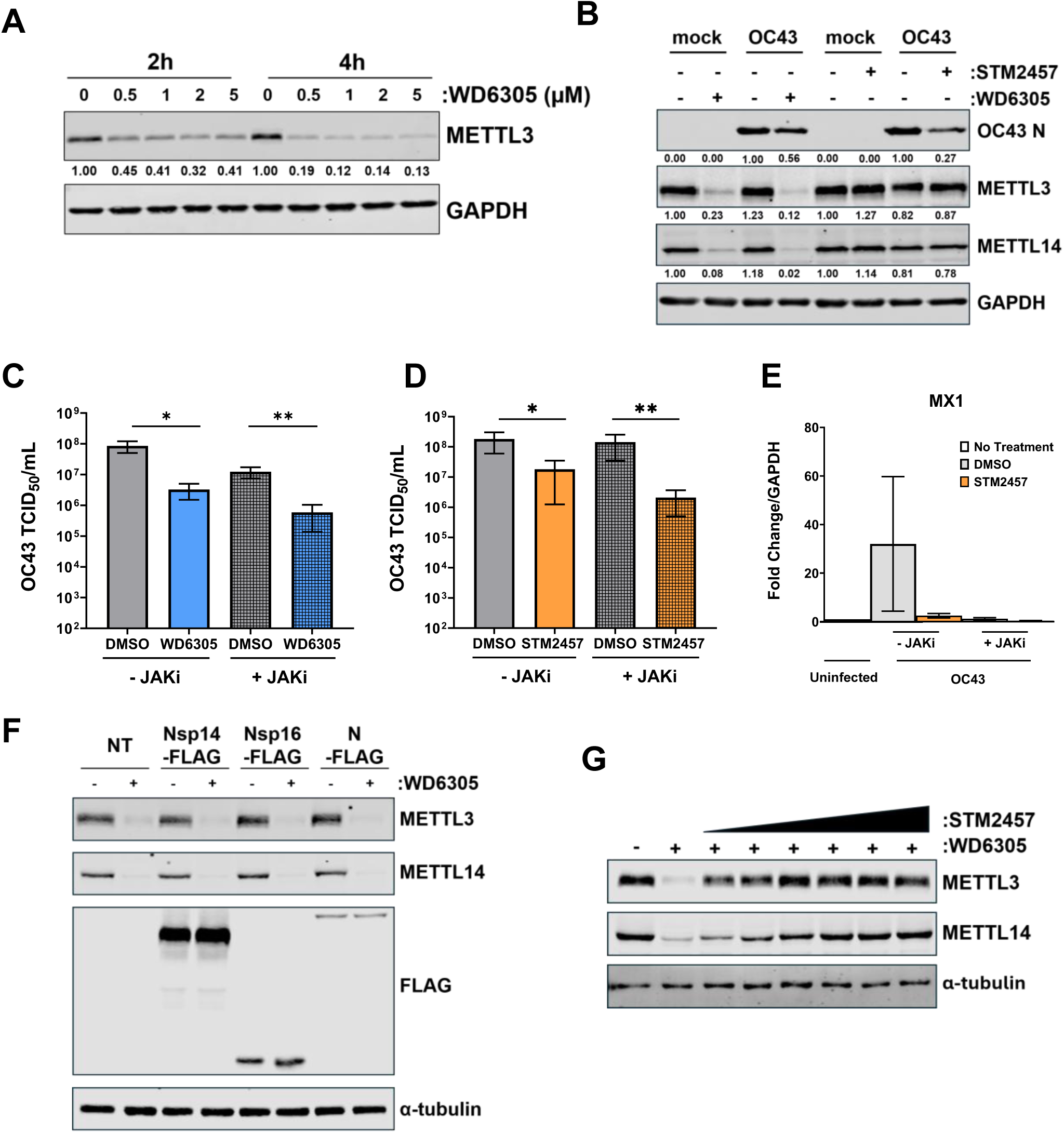
A METTL3-targeting PROTAC has an IFN-independent antiviral activity. (A) MRC-5 cells were treated with WD6305 and immunoblotted for METTL3 and GAPDH. METTL3 abundance normalised to GAPDH and relative to 0 µM for 2h or 4h treatment time is indicated. (B) MRC-5 cells were treated with either WD6305 (1 µM) 4 h prior to and during infection or STM2457 (30 µM) during infection, alongside appropriate DMSO controls and infected at MOI 3. At 24 hpi protein lysates were collected for immunoblotting. Abundance of N, METTL3 and METTL14 normalised to GAPDH and relative to control treatments is indicated. (C, D, E) MRC-5 cells were infected at MOI 0.001 in the presence of WD6305 (1 µM) (C) or STM2457 (30 µM) (D) and JAKi (10 µM) or equivalent concentrations of DMSO. WD6305-treated cells were also treated for 4h prior to infection. Viral supernatants were collected at 48 hpi, analysed by TCID50 assay and titres plotted as mean ± SEM, n ≥ 3. Statistical significance was tested on transformed values by a two-tailed paired t-test. (E) Total RNA was harvested from cells in (D) and subject to RT-qPCR for MX1. Fold change values as mean ± SEM (n=3), relative to GAPDH and normalised to untreated uninfected control were plotted. (F) HEK293T cells were transfected with plasmids encoding OC43 Nsp14, Nsp16 and OC43 N with a C-terminal FLAG tag. After 24h cells were treated with WD6305 (2μM) or DMSO for another 24h before immunoblot analysis. (G) HEK293T cells were treated for 6h with WD6305 (2μM) and DMSO or STM2457 in increasing 2-fold concentrations from 0.9375μM to 30μM prior to immunoblotting.

We then compared the ability of the METTL3 PROTAC (WD6305, 1 µM) to the METTL3 inhibitor (STM2457, 30 µM) to inhibit viral protein expression in a high MOI infection (MOI: 3) at 24 hpi, conditions where we previously observed a strong effect by STM2457 (3). Nucleocapsid protein levels, as measured by immunoblotting, were reduced to 27% of control infected cells by STM2457, and to 56% of control by WD6305 (Fig. 8B). Immunoblotting for METTL3 confirmed depletion in WD6305-treated cells, which was enhanced in OC43 infected cells, potentially reflecting enhanced proteasome activity. Depletion of METTL3 led to a concomitant decrease in its writer complex partner METTL14, consistent with prior reports (51, 52). These results indicate that acute depletion of METTL3 can phenocopy the effect of its inhibition on human coronavirus gene expression.

To establish whether WD6305, like STM2457, also inhibits OC43 replication through an interferon independent mechanism, we tested whether a similar antiviral efficacy was obtained when signalling downstream of the interferon receptor was blocked using pyridone 6, a pan-JAK inhibitor. In JAK inhibitor (JAKi) treated and untreated cells, WD6305 led to a similar and significant reduction in released viral titres (26- and 21-fold respectively) (Fig. 8C), consistent with results obtained with STM2457 (Fig. 8D). The efficacy of JAK inhibitor treatment was validated by RT-qPCR for canonical ISG MX1, whose small induction by multicycle OC43 infection in DMSO-treated cells was almost entirely abrogated by JAKi (Fig. 8E).

Lastly, we considered it possible that STM2457 and WD6305 could inhibit viral replication by non-specifically inhibiting a viral enzyme. S-adenosylmethionine (SAM) serves as a universal methyl donor and both STM2457 and WD6305 are designed to bind the SAM-binding pocket of METTL3 (51, 53). Coronaviruses produce two RNA methyltransferases, Nsp14 and Nsp16, that perform the sequential methylations necessary to cap viral RNAs and each possess a SAM-binding pocket (54). To test whether WD6305 can bind and degrade OC43 Nsp14 and Nsp16 each protein was expressed with a C-terminal FLAG tag from transfected plasmids in HEK293T cells and cells subsequently treated with WD6305 (Fig. 8F). FLAG-tagged OC43 N was used as a control viral protein lacking a SAM-binding site. Immunoblotting demonstrated no reduction in Nsp14, Nsp16 or N protein abundance upon WD6305 treatment. METTL3 and METTL14 were however effectively degraded, and this was not ameliorated by the expression of either viral protein. We also tested whether STM2457 can block access by WD6305 and alleviate METTL3 degradation, confirming a shared binding pocket. Cells were treated for 6 h with WD6305 (2 µM), and an increasing concentration of STM2457, from 0.9 to 30 µM (Fig 8G). Even the lowest concentration of STM2457 impacted METTL3 degradation by WD6305, with METTL3 levels returning those of untreated cells from 3.75 µM STM2457, demonstrating that STM2457 and WD6305 compete to bind the same site on METTL3. Thus, our data indicate that WD6305 binds METTL3 specifically and its antiviral activity is due to depletion of METTL3. Together our results establish that a METTL3-targeting PROTAC has potential application as a host-directed antiviral therapeutic, and, as with METTL3 inhibitor STM2457, exerts an interferon independent antiviral activity.

## Discussion

Several studies have linked the antiviral effects of METTL3 depletion or inhibition during coronavirus infection to induction of innate immune responses. In SARS-CoV-2 infection, for example, the phenotype associated with METTL3 depletion has been attributed in part to enhanced recognition of viral RNA by RIG-I and subsequent activation of innate immune signalling (4). In contrast, the replication of the swine alpha-coronavirus transmissible gastroenteritis virus (TGEV) was recently shown to be restricted by METTL3, and its inhibition by STM2457 was accompanied by decreased ISG expression (7). Even in uninfected cells, METTL3 inhibition can be sufficient to elicit a dsRNA-dependent IFN response (55, 56). Our findings indicate that METTL3 activity can also support coronavirus replication independently of the regulation of innate immune responses. In the OC43 infection model used here, the antiviral activity of STM2457 was not accompanied by increased IFN signalling or broad induction of ISGs and was maintained when host transcription was inhibited. Furthermore, neither RIG-I nor MDA5 were required for the effect of STM2457, and ISGs previously identified as potent restrictors of OC43 were also not required for the phenotype. These observations support the existence of an additional IFN-independent function of METTL3 methyltransferase activity, that contributes directly to coronavirus replication, revealed using a virus that almost completely evades immune activation in a lung fibroblast cell line.

During OC43 infection of MRC-5 cells inhibition of signalling downstream of the interferon receptor did not notably increase OC43 replication, with RIG-I and MDA5 knockdown also leading to only slight increases in viral titres. Consistent with this, and published transcriptome and translatome data sets from MRC5 cells (26), we did not detect significant induction of canonical ISGs during OC43 infection. ISG expression in MRC-5 cells was shown to be highly responsive to IFN treatment however, indicating that OC43 successfully replicates either without detection, or by engaging highly effective countermeasures. While OC43 encodes fewer accessory proteins than the related betacoronavirus SARS-CoV-2, several well characterized immune antagonists are conserved (57). For example, OC43 expresses homologs of non-structural protein 1 (Nsp1), which suppresses host gene expression post-transcriptionally, and Nsp15 (endoU) which suppresses excess dsRNA accumulation, that share these functions (58–60). OC43 NS2 also has 2’,5’-phosphodiesterase activity which prevents RNase L activation by OAS (61). This likely explains why OAS depletion did not affect OC43 titres unless OAS levels were heightened by IFN exposure. OC43 infection of MRC-5 cells thus provides insight into viral biology in the absence of innate immune maelstrom, however whether the antiviral effect of METTL3 inhibition on OC43 could be further augmented by effects on immune sensing and responses in a different cellular context is yet to be determined. Of note, in our hands, infections of A549 and BEAS-2B cells with OC43 yielded similarly muted ISG responses.

We find that the antiviral effect of METTL3 inhibition is maintained even when host transcription is blocked and its effect on viral RNA accumulation can be detected within a few hours. Since m^6^A deposition is believed to occur largely co-transcriptionally, this could be explained by a requirement for METTL3 to act directly on viral RNA and our prior identification of m^6^A on SARS-CoV-2 and OC43 RNAs (3). How this interaction impacts viral processes is unclear. Biochemical studies have highlighted the potential for m^6^A to disrupt RNA:RNA interactions between coronavirus transcription regulatory sequences in the leader and genome body required for the template switching that generates subgenomic RNAs (12). Work with reporter constructs in uninfected cells also suggests that m^6^A in the genomic 5’ UTR could promote translation by influencing RNA structure (13). This could in turn affect the production of RdRP subunits translated directly from ORF1ab of the genomic RNA, impacting viral RNA amplification. It also remains possible that, similar to the regulation of cellular transcripts, m^6^A on viral RNA leads to differential binding of RBPs that impact its stability, translation or structure. Experiments identifying proteins associated with OC43 and SARS-CoV-2 RNA isolated m^6^A reader protein IGF2BP3 (62) and we previously showed that OC43 was regulated by YTHDF family m^6^A reader proteins (3). A challenge for the field is to establish which modified sites are important for coronavirus replication. Mapping with high confidence the sites with highest m^6^A modification stoichiometry on viral RNAs, particularly the 30 kb long genomic RNA, is difficult. Nanopore DRS, arguably the most sensitive method at present, proceeds from the 3’ end, with few reads extending beyond 10 kb. Compounding matters further, sgRNAs dominate the transcriptome of infected cells (26) and alternative short-read based methods are unable to distinguish genomic from sgRNAs. Furthermore, though negative sense coronavirus RNAs contain m^6^A consensus DRACH motifs, they are present at much lower abundance than positive sense RNAs and do not possess a poly(A) tail required for nanopore DRS adaptor ligation. Detection of RNA modifications on these will be technically challenging and require enrichment techniques. Moreover, once frequently modified sites are identified, combinatorial mutations may be required to establish those sites that are most important.

Here we demonstrate that targeting METTL3 using the PROTAC WD6305 has an antiviral effect, reducing both viral titres and viral gene expression. This recapitulated our prior findings with METTL3 knockdown (3), without requiring days long incubation for constitutive protein turnover that can lead to secondary effects on host gene expression. This supports use of METTL3 PROTACs not only as a novel therapeutic modality with applications in other viral infections demonstrated to be METTL3-dependent, such as HSV-1 (63) and IAV (64, 65), but as a research tool to orthogonally validate observations made using METTL3 catalytic inhibitors and probe the non-enzymatic functions of METTL3. Nonetheless, in our experimental conditions we observed a greater effect on viral N protein expression by STM2457-mediated inhibition of METTL3 than WD6305-meditated protein depletion, which could suggest that the small residual pool of METTL3 remaining following PROTAC treatment is sufficient to minimally support viral gene expression. Consistent with this notion, we previously found only a minority of METTL3 is present in the cytoplasm of MRC-5 cells, and that this is unchanged by OC43 infection (3). Whether coronaviruses manipulate METTL3 to facilitate its cytoplasmic activity is an open question.

Overall, our results reveal an interferon-independent mechanism of METTL3 regulation of a human betacoronavirus and highlight a new antiviral therapeutic opportunity by targeting this host enzyme using a PROTAC.

## Acknowledgements

HMB is supported by the Academy of Medical Sciences (SBF008\1027) and the Medical Research Council (MR/Z505523/1). ACW is supported by grants from the National Institute of Allergy and Infectious Diseases (R01-AI176335 and R01-AI170583). DO is supported by the Walter Benjamin Programme of the Deutsche Forschungsgemeinschaft (DFG), project 554758329. We thank Daniel Goncalves-Carneiro for helpful discussions.

